# An overlooked microbial pathway links organic nitrogen turnover in composts to nitrous oxide formation

**DOI:** 10.64898/2026.07.12.738090

**Authors:** Ju Yong Lee, Minho Lee, Sojung Yoon, Min Joon Song, Sukhwan Yoon

## Abstract

Biological N_2_O production from organic nitrogen is generally assumed to require canonical nitrification, which generates oxidized nitrogen that subsequently fuel denitrification. Whether this paradigm universally applies to nitrogen-rich microbial communities remains unclear. Here, we investigated N_2_O production across an industrial poultry manure composting process and found that substantial N_2_O formation occurred despite the apparent absence of canonical ammonia oxidation. Neither allylthiourea inhibition nor metagenomic analyses provided evidence for ammonia-oxidizing microorganisms or their activity. Instead, metagenomic analyses identified abundant bacterial nitric oxide synthase (*bNos*) genes, many of which were phylogenetically affiliated with Bacilli, the dominant bacterial group throughout composting. Physiological experiments with *Bacillus* isolates demonstrated a nitrification-independent route in which L-arginine was oxidized to NO_2_⁻/NO_3_⁻, consistent with bNOS-mediated NO formation followed by abiotic oxidation. Recovery of ^15^N-labelled N_2_O following ^15^NO_2_⁻ addition established NO_2_⁻ as an immediate precursor of aerobically produced N_2_O, confirming that the oxidized nitrogen generated through this alternative route subsequently fueled denitrification. Metagenomic analyses further revealed extensive denitrification potential but comparatively low *nosZ* abundance. Together, these findings identify a previously overlooked route linking organic nitrogen turnover to denitrification independently of canonical nitrification, thereby expanding current models of microbial N_2_O production in composts and potentially other protein-rich thermophilic environments.

## Introduction

Nitrous oxide (N_2_O) is a potent greenhouse gas with a global warming potential 298 times that of CO_2_ and contributes approximately 5% of the net anthropogenic radiative forcing^1^. N_2_O is produced primarily through biological processes, with nitrification and denitrification recognized as its principal sources^2, 3^. In the canonical N_2_O-centric view of the nitrogen cycle, organic nitrogen, the dominant nitrogen pool in terrestrial ecosystems, is first mineralized to NH_4_^+^/NH_3_ and subsequently aerobically oxidized through nitrification. This process directly produces N_2_O while generating NO_2_⁻ and NO_3_⁻ that subsequently fuel denitrification and additional N_2_O production^2, 3^. As such, biological N_2_O formation from organic nitrogen is generally assumed to require nitrification^3, 4, 5, 6^.

Composting of organic wastes represents an important source of anthropogenic N_2_O emissions and has therefore served as a model system for investigating microbial N_2_O formation^3, 4, 5, 7, 8^.Current models generally interpret N_2_O emissions from manure composting within the canonical framework in which mineralization converts organic nitrogen to NH₄⁺, followed by sequential nitrification and denitrification, both of which contribute to N_2_O production^2, 3, 4, 5, 6^. These processes are typically considered to occupy distinct positions along the oxic-anoxic continuum, with nitrification requiring oxygen and denitrification proceeding primarily under oxygen limitation^2, 9^. Nevertheless, the intrinsic spatial heterogeneity of composting manure permits the simultaneous occurrence of these N_2_O-producing nitrogen transformation processes^8, 10^.

Despite this prevailing paradigm, whether canonical ammonia oxidation is a universal prerequisite for biological N_2_O formation from organic nitrogen deserves critical re-examination. In particular, little attention has been paid to whether ammonia-oxidizing microorganisms are consistently present and active in nitrogen-rich composting systems that experience elevated temperatures and physicochemical conditions potentially unfavorable for canonical nitrifiers^5, 6^. Nevertheless, substantial N_2_O emissions have repeatedly been reported from animal manure composts, raising the possibility that organic nitrogen may fuel biological N_2_O production through alternative microbial pathways that bypass canonical ammonia oxidation^3, 4, 5, 7^. Such a mechanism would require a substantial revision or expansion of current conceptual models of biological N_2_O formation.

To address this question, we investigated the microbial mechanisms underlying N_2_O production in poultry manure and compost samples collected across successive stages of industrial composting. We found that canonical ammonia oxidation was not required for sustained biological N_2_O production. Instead, our results support a previously unrecognized nitrification-independent pathway in which organic nitrogen is converted to NO_2_⁻ and NO_3_⁻, thereby supporting downstream N_2_O formation. By integrating pure-culture physiological experiments with ^15^N isotope tracing in manure incubations, we reconstructed a microbial pathway linking organic nitrogen mineralization to N_2_O production independently of canonical ammonia oxidation. Together, these findings expand the current conceptual framework of biological N_2_O formation.

## Methods

### Composting site description and manure sampling

The manure and compost samples used in this research were collected in August 2024 from a composting factory located in Sancheong, South Korea (35°22’11.2“N, 127°59’08.1“E). The facility processes, per day, approximately 9 tons (wet weight) of poultry waste collected from nearby chicken farms into approximately 10 tons of fertilizer for distribution to local farmlands. Incoming wastes are mixed with sawdust (approximately 10% v/v) and a commercial “effective microorganisms” product of undisclosed composition, followed by two weeks of continuous (uninterrupted) mechanical mixing. After aerobic composting, the mixture undergoes six months of curing in an undisturbed pile. Subsequently, the final fertilizer pellets are produced by blending the composted mixture with air-dried fresh manure at a mass ratio of 3:1. The process is illustrated in Fig. 1a.

**Fig. 1.**
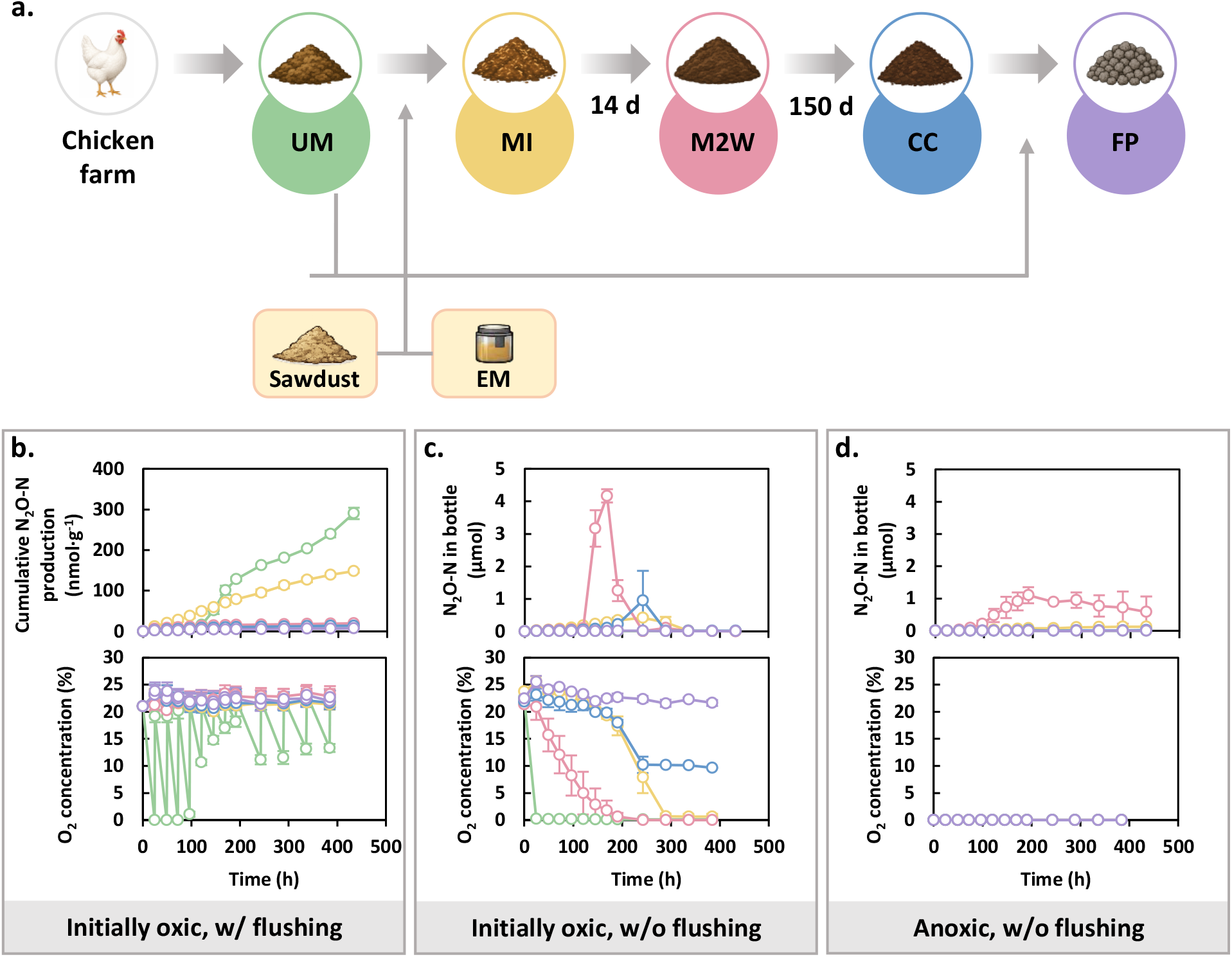
N_2_O production of poultry manure and composts under contrasting headspace regimes. (a) Schematic overview of the poultry manure composting process. UM, untreated manure; MI, manure mixed with sawdust and effective microorganisms; M2W, mixture after two weeks of composting; CC, cured compost, FP, finalized fertilizer pellets. (b-d) N_2_O production by manure and composts sampled at different composting stages. (b) Cumulative N_2_O−N production under ambient air with headspace replenishment every 8 h. (c) Time-course of N_2_O−N in bottles under initially oxic conditions without headspace replenishment. (d) Time-course of N_2_O−N in bottles under an N_2_ headspace. Headspace O_2_ concentrations were also monitored throughout incubations. Error bars indicate standard deviations (*n* = 3). Additional data from these experiment, including two-point measurements of inorganic nitrogen concentrations, are provided in Supplementary Figure 1.

The samples for chemical characterization, incubation experiments, and microbiome characterization were collected at five different steps during this process: 1) untreated raw manure (UM), 2) manure mixed with sawdust and effective microorganisms (MI), 3) mixture after two weeks of composting (M2W), 4) cured compost (CC), and 5) finalized fertilizer pellet (FP). Additionally, sawdust was collected for characterization of the microbial community composition. Each set of samples was collected from three different locations within the surface layer (0–2 cm depth) of the manure/compost pile and combined into a single composite sample. After removing visible impurities, the manure and compost samples were transferred into pre-sterilized conical tubes and transported to the laboratory in a cooler filled with ice. The samples for incubation experiments were stored at 4℃ for no longer than 14 days. Aliquots of the samples for chemical and sequencing analyses were stored at −20℃ and at −80℃, respectively.

### Measurements of N_2_O emissions from manure/compost samples

The manure and compost samples were incubated without any amendment, to approximate *in*-*situ* N_2_O emission at the composting facility. Three grams (wet weight) of each sample were added to an autoclaved 140-mL glass bottle, which was then sealed with a bromobutyl stopper fitted into a GL45 open-topped screw cap. The bottles were initially flushed with either air (>99.999%; Jeilgas Co., Sejong, South Korea) or N_2_ (>99.999%; Jeilgas Co., Sejong, South Korea). Each sample was incubated under three headspace conditions: (i) N_2_ headspace without replenishment (anoxic), (ii) air headspace without replenishment, and (iii) air headspace with periodic replenishment every 24 h. N_2_O concentrations were measured at 24-h intervals over 18 days. For condition (iii), N_2_O concentration was measured immediately before each headspace replenishment. Incubations were performed at 25℃ to approximate conditions in the surface layers of the compost pile, where temperatures are typically closer to ambient air and gas exchange with the atmosphere is less likely to be limited. All incubation experiments were performed in triplicates, in the dark, without agitation.

### Nitrogen turnover and N_2_O production potential by native microbial communities in manure and composts

Aerobic and anaerobic microbial nitrogen turnovers were also examined in closed-batch suspended cultures prepared with the manure/compost samples (UM, MI, M2W, CC, and FP) as inocula. Cultures were prepared in 160-mL serum bottles containing 60 mL of 10 mM phosphate buffer solution prepared with Na_2_HPO_4_ and KH_2_PO_4_ and adjusted to pH 7.2. After autoclaving the bottles, 3 g (wet weight) of the manure/compost sample was suspended to the medium. The bottles were then sealed with butyl rubber stoppers and aluminum crimps. Aerobic incubation was started with air in the headspace, and flushed every 8 hours with compressed air. To a subset of the aerobic cultures, 100 µM allylthiourea (ATU) was added to inhibit ammonia oxidation.^33^ Anaerobic cultures were prepared by bubbling >99.999% N_2_ gas for 20 minutes after sealing the bottles. Acetylene gas (>99.99%; Jeilgas Co., Sejong, South Korea) was added to a subset of the anoxic cultures at a concentration of 10% v/v in the headspace to estimate potential denitrification activity by inhibiting N_2_O reduction to N_2_.^34^ The NO_3_^−^, NO_2_^−^, and NH_4_^+^ concentrations in the aqueous phase and N_2_O concentration in the headspace were monitored until no further change to N_2_O concentration was observed.

### Microbial community composition analyses

The manure and compost samples collected from the five stages of poultry waste processing, and also, effective microorganism inoculum and sawdust, were analyzed for their microbial community compositions. Genomic DNA was extracted using the DNeasy PowerSoil Pro Kit (Qiagen, Hilden, Germany). The V3–V4 hypervariable region of the 16S rRNA gene was amplified with 341F (5’-CCTACGGGNGGCWGCAG-3’) and 805R (5’-GACTACHVGGGTATCTAATCC-3’) primer pair, and the resulting amplicons were sequenced on an Illumina MiSeq platform (San Diego, CA). The sequence data were processed using QIIME2 v2024.10.^35^ Briefly, raw paired-end sequences were trimmed using Trimmomatic v0.39 and merged using the ‘denoise-paired’ plugin.^36, 37^ The amplicon sequence variants (ASVs) were taxonomically classified using the Naive Bayes classifier, trained on the SILVA SSU Ref NR 99 database (release 138.2), with a confidence threshold of 0.8.^38^ The ASVs that could not be unambiguously assigned genus-level taxa, or lacked identified genus-level relative, were clustered into operational taxonomic units (OTUs) at 97% identity threshold and reported at the lowest confidently identifiable taxonomic rank.^39^

### Gene-centric metagenomic analyses

The extracted DNA samples were subjected to shotgun metagenomic sequencing to identify and quantify functional genes relevant to nitrogen turnover and N_2_O emissions. Sequencing was performed on an Illumina NovaSeq 6000 platform, generating approximately 10 Gb of paired-end reads (2×150 bp) per sample. Raw 16S rRNA gene amplicon and shotgun metagenomic sequencing data have been deposited in the NCBI Sequence Read Archive (SRA) under BioProject accession number PRJNA1483828. The raw reads were quality-filtered and trimmed using fastp v0.24.0 with parameters set to: qualified_quality_phred: 20 and length_required: 70.^40^ *De novo* assembly was performed using MetaSPAdes v3.15.3 with a minimum contig length threshold of 2000 bp.^41^

Contigs were analyzed without binning. Gene-coding sequences were predicted and translated *in silico* using Prodigal v2.6.3 and annotated using Kofamscan v.1.3.0.^42, 43^ Resulting annotations were cross-validated using DIAMOND-BLASTX v2.1.10 against the NCBI nr protein database (accessed on January 16, 2025), applying an E-value cutoff of 1E–10.^44^ The primary target genes included *amoA*, *hao*, *napA*, *narG/nxrA* (distinguished based on phylogenetic context), *nirS*, *nirK*, *norB*, and *nosZ* representing ammonia oxidation and denitrification pathways. Kofamscan-annotated *norB* were further classified as qNOR or cNOR using heme-copper oxidase (HCO) family hidden Markov models, and *nosZ* were categorized into clade I and II using InterProScan v.5.73-104.0 with reference to the following databases: NCBIfam, Pfam, TIGRFAM.^45, 46^ Additionally, NO-related genes including *nos*, *hmp*, *norV*, and *nsrR* were examined. Relative abundances of the predicted coding sequences were determined by mapping quality-trimmed raw reads onto the coding sequences using BWA v0.7.18-r1243-dirty with default parameters.^47^ Duplicate reads were removed using the ‘markdup’ command in SAMtools v1.21.^48^ Gene abundances were estimated as RPKM (reads per kilobase per million mapped reads) values, calculated using the ‘genomecov’ command in BEDtools v2.31.1.^49^

Taxonomic compositions of *narG*, *nirK*, *norB*, *nosZ*, and *nos* genes were analyzed with the predicted coding sequences and relative abundances quantified using corresponding RPKM values. For each target gene, predicted full-length amino acid sequences were combined with reference sequences downloaded from KO_list (accessed January 20, 2025). These sequences were clustered using CD-HIT v4.8.1 with the amino acid-level similarity thresholds of 75%, 75%, 85%, 90%, and 90% for *narG*, *nirK*, *norB*, *nosZ*, and *nos*, respectively. For each cluster, a representative sequence was selected based on sequence coverage, and phylogenetic trees were constructed using these representative sequences. Cluster abundance was calculated as the sum of *rpoB*-normalized coverages of the metagenomic contigs encoding genes assigned to each cluster.

### NO_2_^−^ production from L-arginine in two *Bacillus* isolates

*Bacillus subtilis* strain ATCC 6051a (acquired from the American Type Culture Collection, ATCC) and *B. licheniformis* strain BCRC 11702 (acquired from Korean Collection for Type Cultures, KCTC), both with fully sequenced genomes (GCF_003417435.1 and GCF_034478925.1, respectively), were examined for the ability to metabolize L-arginine to N_2_O or its potential precursors (NO, NO_2_^−^, and/or NO_3_^−^). The two strains were selected based on their nitrogen functional gene inventory, as they harbor a putative bacterial nitric oxide synthase gene (*bNos*), while lacking *nirK* and *nosZ* (Supplementary Table 9). A modified MR-1 medium was prepared by dissolving 30 g NaCl, 0.82 g Na_2_HPO_4_, 0.53 g KH_2_PO_4_, and 1 mL of a 1000X trace metal solution (Supplementary Table 10) in 1 L of Milli-Q water. After autoclaving, 5 mL of a 200X vitamin stock solution (Supplementary Table 10) was aseptically added to the medium, which was then dispensed to presterilized 160 mL serum bottles in 100-mL aliquots. Precultures were grown with 20 mM acetate as carbon and non-fermentable electron source and 10 mM NOs_3_^−^ and 5 mM NHs_4_^+^ as nitrogen source. The bottles were sealed with ambient air in the headspace, which was flushed with air when O_2_ concentration dropped below 0.15%. The precultures were harvested at OD_600nm_∼0.058, washed with 10 mM phosphate buffer, and resuspended in 10 mL medium. To a set of triplicate cultures, filter-sterilized L-arginine stock solution (stored at 4℃) was added to a concentration of 0.5 and 1 mM. NH_4_^+^ was supplemented at 0.1 mM as an additional nitrogen source in case the *Bacillus* isolates are unable to metabolize L-arginine. Control cultures were prepared identically, but without L-arginine addition. All experiments with the *Bacillus* isolates were performed at 30℃.

### Verification of N_2_O production from ^15^NO_2_^−^ in oxic compost suspension

To verify that NO_2_^−^ could be further reduced to N_2_O in oxic composts, aerobic suspensions of M2W samples were incubated with ^15^NO_2_^−^, and N_2_O production was monitored. Incubations were performed in 26-mL glass tubes containing 16 mL of 10 mM phosphate buffer (pH 7.2) and 0.1 g (wet weight) M2W compost. After Na^15^NO_2_ was added to 1 mM, tubes were sealed and incubated aerobically, with the headspace flushed every 8 h with a stream of compressed air. Headspace samples for ^46^N_2_O measurements were withdrawn immediately before headspace flushing. Abiotic controls with autoclaved M2W compost were examined in parallel.

### Analytical methods

The physicochemical properties of the chicken manure and compost samples were characterized immediately after collection. The pH was measured by suspending 1 g of the sample in 5 mL of Milli-Q water. Total nitrogen (TN) was determined using the standard Kjeldahl titration method.^50^ Total organic carbon (TOC) was determined using the Walkley-Black wet oxidation method by reacting 0.5 g of sample with 20 mL of sulfuric acid (95–98%, w/w) and 10 mL of 1 N_2_K_2_Cr_2_O_7_.^51^ For quantification of NH_4_^+^, NO_2_^−^, and NO_3_^−^, 1 g (wet weight) of sample was suspended in 5 mL of 2 M KCl solution and agitated for one hour at 200 rpm. The suspension was then allowed to settle for 30 minutes, and the supernatant was filtered through a 0.22 µm membrane filter (Advantec MFS, Inc., Tokyo, Japan). The filtrate was analyzed using previously described colorimetric assays.^52, 53, 54^ N_2_O concentrations, including unlabeled N_2_O, ^45^N_2_O, and ^46^N_2_O, were monitored using a QP2010 gas chromatography-mass spectrometry system (Shimadzu, Kyoto, Japan) equipped with an RT-Q-bond column (30 m x 0.32 mm, 10 µm film thickness).^52, 55^ Calibration standards for ^46^N_2_O were generated biologically using *Pseudomonas stutzeri* strain DCP-Ps1 from Na^15^NO_2_. Briefly, 0.1 mL of strain DCP-Ps1 culture grown with 10 mM sodium acetate and 1 mM KNO_3_^−^ under N_2_ headspace was transferred into sealed 26-mL glass tubes containing 16 mL modified MR-1 medium supplemented with concentrations of Na^15^NO_2_ ranging from 0 to 0.1 mM^56^. The headspace was replaced with He (>99.999%; Jeilgas Co., Sejong, South Korea), and acetylene was added to inhibit N_2_O reduction, allowing ^46^N_2_O to accumulate as the terminal denitrification product^34^. ^45^N_2_O standards were generated chemically using Na^15^NO_2_ and Na^14^N_3_.^57^ Briefly, Na^15^NO_2_ (0 to 0.1 mM) was reacted with 2 M Na^14^N_3_ at pH 4.5 under a He headspace, producing ^45^N_2_O through acid-catalyzed nitrosation of azide by HNO_2_ derived from Na^15^NO_2_. The total amount of N_2_O in a container was calculated from headspace concentrations assuming gas-liquid equilibrium, using a dimensionless Henry’s constant (gaseous/aqueous) of 1.68 at 25℃.^58^ L-arginine concentration was quantified using a Prominence high performance liquid chromatograph system (Shimadzu, Kyoto, Japan), equipped with a reverse-phase C18 column.^59^

### Data availability

The 16S rRNA amplicon and metagenomic sequencing data generated in this study have been deposited in the NCBI Sequence Read Archive under BioProject accession number PRJNA1483828. Detailed information on the metagenomic analyses is provided in Supplementary Tables 3–8. All relevant data are included in the paper and its source data files. Source data are provided with this paper.

## Results

### Reduced nitrogen dominates the compost nitrogen pool

Five samples representing successive stages of an industrial poultry manure composting process were analyzed: untreated manure (UM), manure immediately after mixing with bulking agent and microbial inoculum (MI), 2-week compost (M2W), cured compost (CC), and finished pellets (FP; properties summarized in Supplementary Table 1). All manure and compost samples were nitrogen-rich, with total nitrogen (TN) exceeding 1% (wet weight). Based on TN and inorganic nitrogen measurements, >99.8% of nitrogen was present in reduced forms, i.e., organic nitrogen and NH_4_⁺, despite prolonged exposure to atmospheric oxygen. Gravimetric water content varied substantially among samples, ranging from 16.0% in FP to 68.2% in UM, suggesting differences in oxygen diffusion and the potential formation of localized low-oxygen microsites.

### Substantial N_2_O production occurs across oxic and anoxic conditions

All manure and compost samples produced N_2_O under each of the three incubation regimes, although the magnitude and temporal dynamics varied markedly (Fig. 1, Supplementary Fig. 1). During continuously oxic incubation maintained by periodic headspace replacement, cumulative N_2_O production ranged from 7.3±0.1 to 290±14 nmol·(g wet weight)^−1^, with the highest production observed in the UM sample (Fig. 1b). In the UM, MI, and M2W samples, cumulative N_2_O production represented 0.035, 0.009, and 0.001% of the initial TN, respectively.

When headspace oxygen was not replenished, transient N_2_O accumulation was observed in all samples except FP, consistently peaking as oxygen concentrations declined (Fig. 1c). The highest peak was observed in the M2W sample (4.2±0.2 µmol N_2_O-N) 169 h after incubation began, when headspace O_2_ had declined to 1.7%. Pronounced transient peaks were also observed in the MI and CC samples after O_2_ concentrations decreased to approximately 10%. Under anoxic conditions, substantial N_2_O production occurred only in the M2W sample, reaching a maximum of 1.1±0.2 µmol N_2_O-N after 191 h (Fig. 1d). In all unreplenished incubations exhibiting measurable N_2_O production, N_2_O concentrations declined after reaching a peak, indicating active biological N_2_O reduction under oxygen-limited or anoxic conditions.

Changes in inorganic nitrogen pools suggested that N_2_O production could not be explained solely by reduction of pre-existing oxidized nitrogen pool. For example, in the M2W sample incubated under initially oxic condition without headspace replenishment, the peak N_2_O-N amount (4.2±0.2 µmol at 169 h) exceeded the net decrease in the combined NO_3_⁻ and NO_2_⁻ pool (<1.0 µmol, Supplementary Fig. 1b). Moreover, multiple samples showed an increase in the combined NO_2_^−^ and NO_3_^−^ pool. Together, these observations indicate that oxidized nitrogen was regenerated from reduced nitrogen during incubation.

### N_2_O production in composts is independent of canonical nitrification

To determine whether canonical ammonia oxidation contributed to N_2_O production, suspended incubations of the MI, M2W, and CC samples were performed in the presence and absence of the ammonia monooxygenase inhibitor allylthiourea (ATU) (Fig. 2a). Under continuously oxic conditions, with headspace replenished every 8 h to maintain dissolved O_2_ concentrations above 4.3 mg L^−1^, the MI, M2W, and CC suspensions produced 44±2, 222±3, and 27±1 nmol N_2_O-N, respectively. Addition of ATU did not significantly affect N_2_O production, and NO_2_⁻ and NO_3_⁻ dynamics remained indistinguishable from those in untreated controls, indicating that canonical ammonia oxidation contributed little, if at all, to N_2_O production under these conditions.

**Fig. 2.**
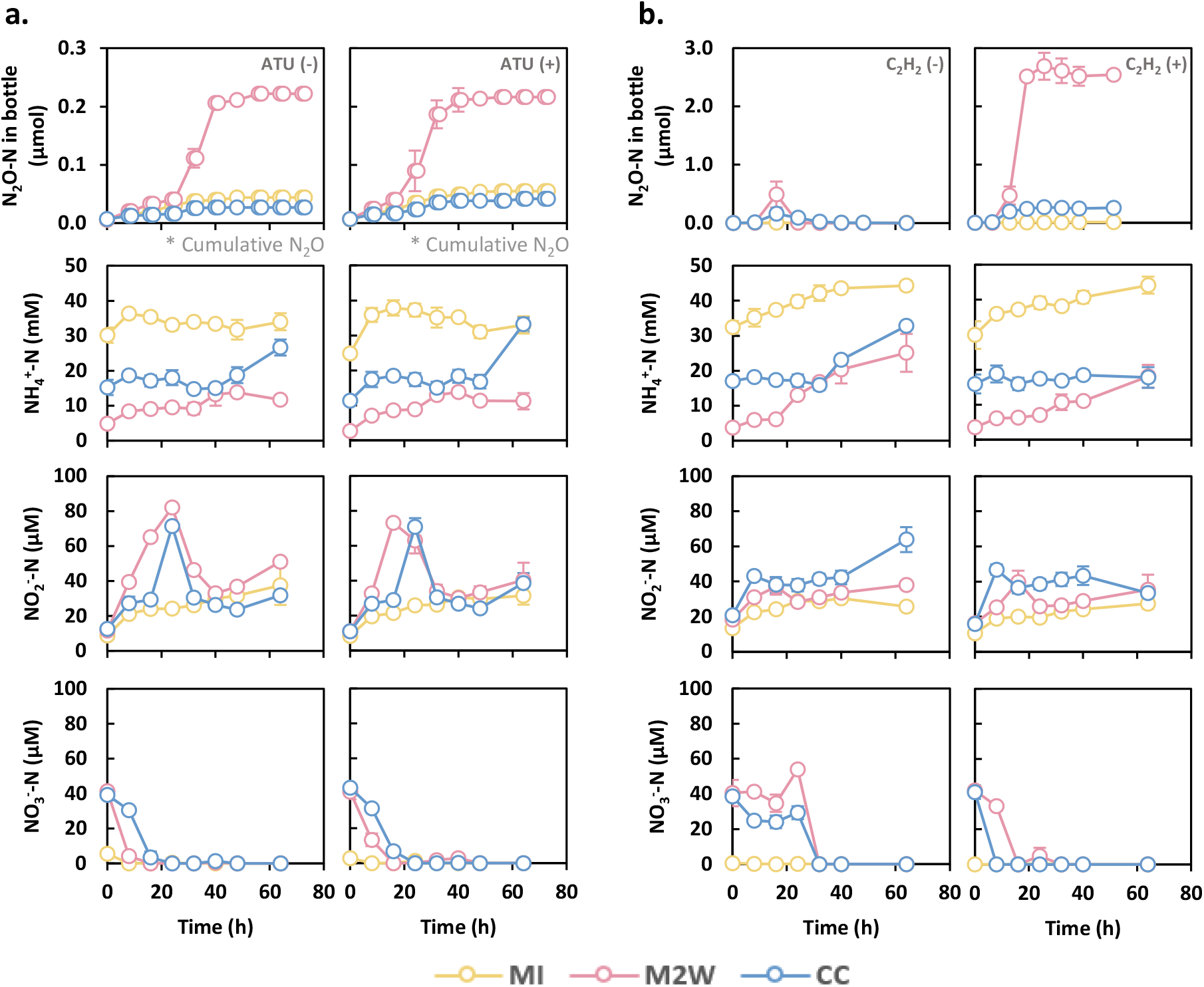
Effects of ammonia oxidation and N_2_O reduction inhibitors on N_2_O production and inorganic nitrogen dynamics in compost suspensions. Compost suspensions prepared from three composting-stage samples (MI, M2W, and CC) were incubated (a) under continuously oxic conditions with headspace replenishment every 8 h in the presence or absence of ATU and (b) under an N_2_ headspace in the presence or absence of acetylene. Cumulative N_2_O-N production is shown in (a), whereas N_2_O-N amounts are shown in (b). NH_4_^+^-N, NO_2_^−^-N, and NO_3_^−^-N concentrations were monitored throughout the incubations. Error bars indicate standard deviations (*n* = 3).

In all three compost suspensions, NO_3_⁻ (initially 4.1, 41.1, and 41.3 µM in MI, M2W, and CC, respectively) was depleted within 24 h, coincident with accumulation of NO_2_⁻. However, NO_2_⁻ production exceeded NO_3_⁻ consumption by at least 1.7-fold and continued after complete depletion of NO_3_⁻, demonstrating that NO_2_⁻ production could not be explained solely by NO_3_⁻ reduction. Peak N_2_O production in the MI and M2W suspensions (24–32 h) coincided with rapid NO_2_⁻ disappearance, identifying NO_2_⁻ as the most likely immediate precursor of N_2_O.

To determine whether denitrification was responsible for converting the regenerated oxidized nitrogen to N_2_O, anoxic incubations were performed with and without the N_2_O reductase inhibitor acetylene (Fig. 2b). Whereas the M2W and CC suspensions transiently accumulated only 490±222 and 157±13 nmol N_2_O–N, respectively, in the absence of acetylene, inhibition of N_2_O reduction resulted in permanent accumulation of 2,690±230 and 268±34 nmol N_2_O-N. In contrast, the MI suspension produced no detectable N_2_O despite containing 11.9±1.4 µM NO_2_⁻, indicating that active denitrification *sensu stricto* was not established in this sample under anoxic conditions. In acetylene-amended incubations, N_2_O production plateaued within 24 h despite persistently elevated NO_2_⁻ concentrations (>38.6 µM), suggesting that denitrification activity was largely confined to the early stages of incubation even though both electron donor and substrate remained available.

Collectively, these observations indicate that substantial N_2_O production occurred independently of canonical ammonia oxidation, with denitrification as the probable principal pathway converting oxidized nitrogen to N_2_O.

### Compost microbiomes lack canonical ammonia oxidizers and are dominated by Bacilli

Amplicon sequencing revealed pronounced enrichment of Bacilli throughout composting (Fig. 3; Supplementary Tables 2–4). Members of this class accounted for 21.2–87.2% of the microbial communities and increased from 43.6% in untreated manure to 56.6% immediately following addition of sawdust and the commercial “effective microorganisms” inoculum, suggesting that the inoculum itself was largely composed of Bacilli, including *Oceanobacillus*, *Sinibacillus*, and *Gracilibacillus*. Composting process further enriched Bacilli to 78.5% and 88.5% in the M2W and CC samples.

**Fig. 3.**
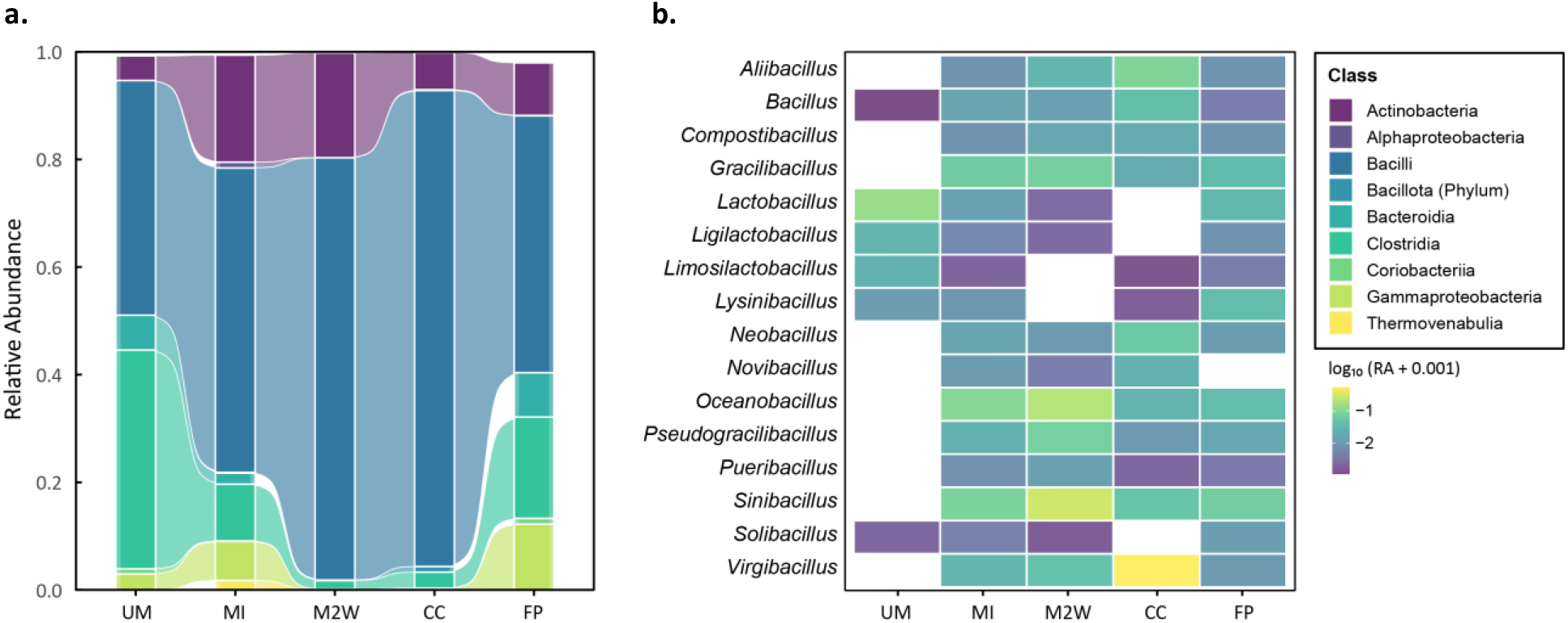
Microbial community shifts across composting stages. (a) Stacked area plots showing class-level microbial community succession across composting stages. (b) Heatmap showing the relative abundance of dominant Bacilli-affiliated genera across composting stages.

No amplicon sequence variants affiliated with archaeal or bacterial ammonia oxidizers or *Nitrospira* were detected in any manure or compost sample (Supplementary Tables 2–4). Other organisms previously reported to convert NH_4_^+^ to NO_2_^−^ were likewise rare. *Methylocystis* spp. accounted for only 0.08% and 0.16% of the MI and CC communities, respectively, whereas *Alcaligenes* spp. were detected only in the finished pellets at 0.03% relative abundance. Thus, microorganisms capable of oxidizing NH_4_⁺ to NO_2_⁻ by previously recognized mechanisms were either absent or detected only at trace abundance throughout the composting process.

Together with the inhibitor experiments, these community data provide independent evidence that canonical ammonia oxidation was unlikely to account for the observed NO_2_⁻ regeneration and N_2_O production, motivating further investigation of alternative microbial pathways capable of generating oxidized nitrogen from organic substrates.

### Metagenomic analyses identify a plausible alternative source of oxidized nitrogen

As canonical ammonia oxidizers were neither detected by community profiling nor inhibitor experiments, shotgun metagenomics was performed to identify alternative microbial pathways capable of regenerating oxidized nitrogen (Fig. 4; Supplementary Tables 5–7). Consistent with the 16S rRNA gene amplicon sequencing analyses, none of the 292.2 million quality-filtered shotgun reads was assigned to known ammonia-oxidizing microorganisms, while sequences assigned to aerobic methanotrophs or the putative heterotrophic nitrifier *Alcaligenes* remained exceedingly rare (<0.01%). Likewise, although *narG*/*nxrA* homologs were abundant (0.07–0.73 copies/*rpoB*), none was affiliated with recognized nitrite-oxidizing bacteria. Collectively, these independent metagenomic observations further exclude canonical nitrification as the primary source of oxidized nitrogen in the composts.

**Fig. 4.**
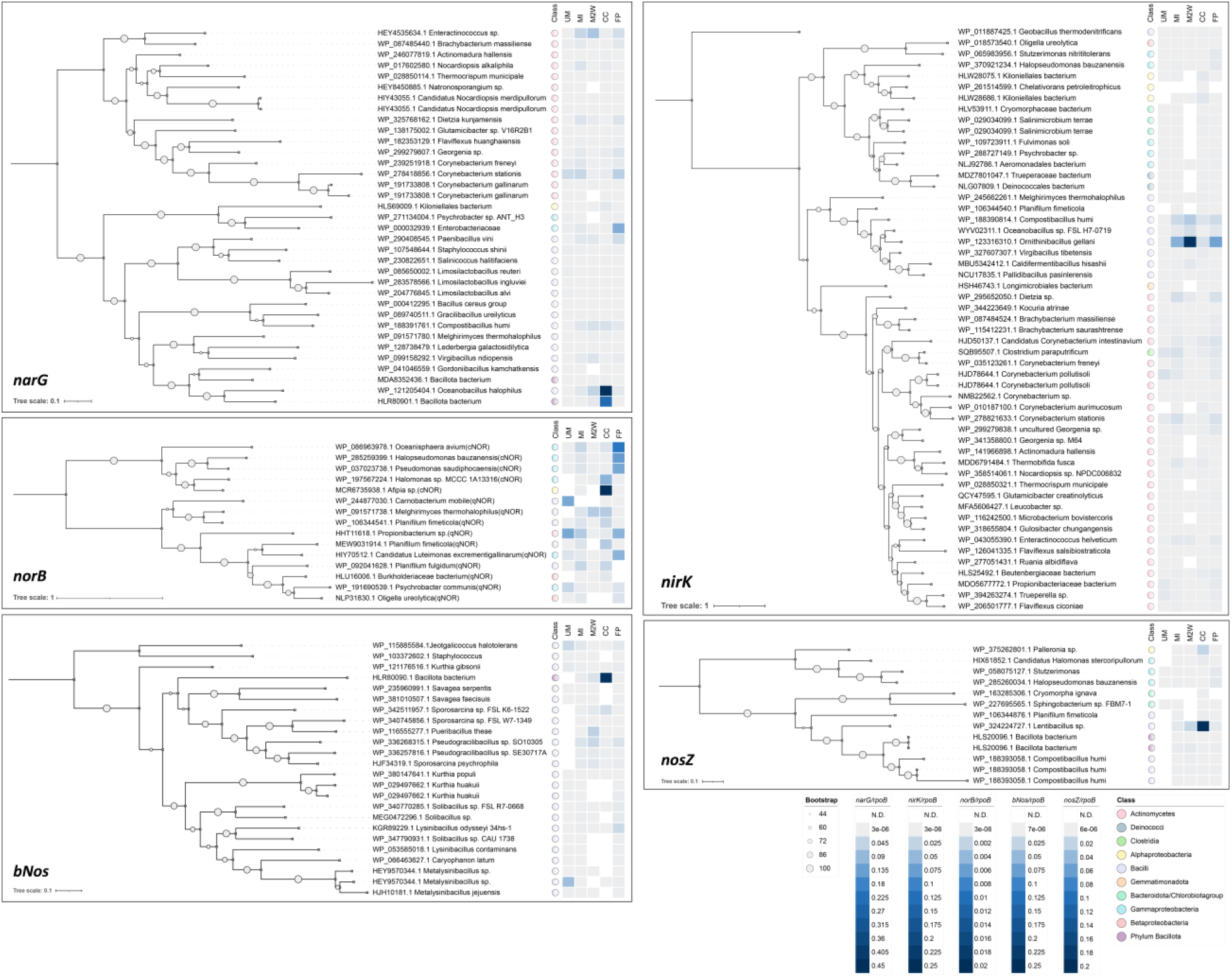
Gene-centric metagenomic analysis of denitrification-related genes across poultry-manure composting stages. Maximum-likelihood phylogenies were constructed for *narG*, *nirK*, *norB*, *nosZ,* and *bNos* genes recovered from metagenomic assemblies. Taxon names correspond to the reference sequences with which the metagenomic genes were clustered. The sizes of the circles at internal nodes indicate bootstrap support values calculated from 1000 replicates. Colored circles adjacent to the representative taxon names indicate class-level taxonomic affiliations. Heatmaps adjacent to each tree display the relative abundance of each gene across the five composting stages (UM, MI, M2W, CC, and FP), expressed as copies per *rpoB*.

Instead, metagenomic analyses identified bacterial nitric oxide synthase (*bNos*) as a potential alternative source of oxidized nitrogen. *bNos* genes were detected in all five metagenomes at abundances exceeding 0.04 copies/*rpoB* and were particularly abundant in the CC sample (0.26 copies/*rpoB*). All recovered *bNos* sequences were affiliated with Bacilli, consistent with the taxonomic composition revealed by amplicon sequencing.

### L-Arginine metabolism enables a nitrification-independent pathway to oxidized nitrogen

To determine whether L-arginine metabolism could generate oxidized nitrogen, and consequently N_2_O, independently of canonical ammonia oxidation, two complementary experiments were performed. First, two *Bacillus* isolates carrying *bNos* were examined for NO_2_⁻ and NO_3_⁻ production from L-arginine (Fig. 5a–e). In *Bacillus licheniformis* strain BCRC11702, amendment with 200 µmol-N L-arginine resulted in transient accumulation of NO_2_⁻ and NO_3_⁻, reaching maxima of 3.43±0.20 and 3.79±0.96 µmol-N, respectively, as approximately 50 µmol-N of added L-arginine was metabolized. Under identical conditions, *Bacillus subtilis* strain ATCC6051a also produced substantial amounts of NO_2_⁻ and NO_3_⁻, reaching maxima of 2.23±0.56 and 12.19±1.59 µmol-N during complete metabolism of the 200 µmol-N L-arginine supplied. Neither uninoculated controls nor cultures lacking L-arginine produced detectable NO_2_⁻ or NO_3_⁻, demonstrating that oxidized nitrogen originated from microbial L-arginine metabolism. Although NO was not detected in the headspace, its rapid abiotic oxidation under oxic conditions likely prevented direct detection.

**Fig. 5.**
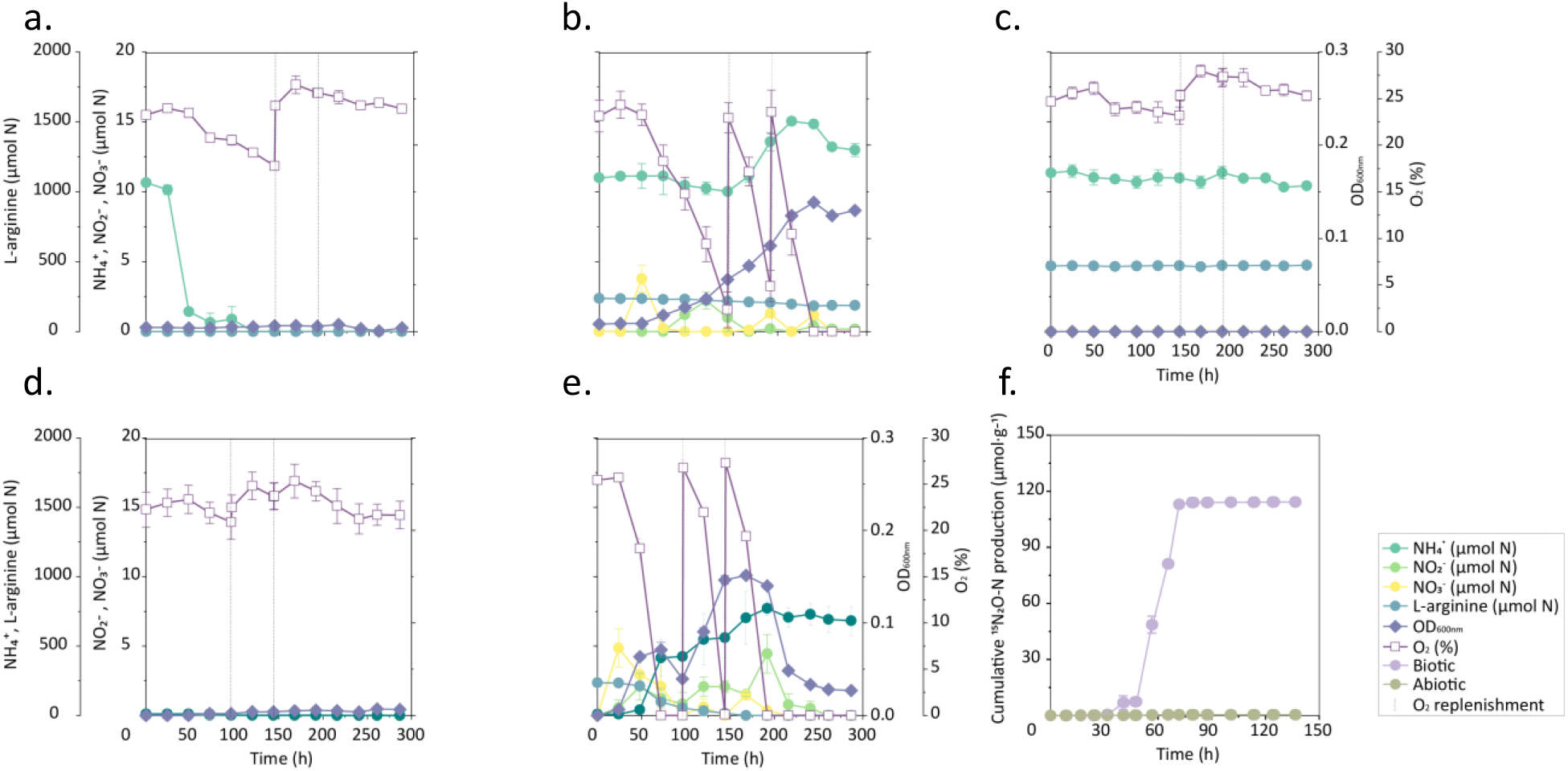
Physiological verification of L-arginine-derived oxidized nitrogen regeneration and subsequent N_2_O production. Time-course measurements of nitrogen species during incubation of *Bacillus licheniformis* strain BCRC 11702 (a, b) and *Bacillus subtilis* strain ATCC 6051a (c, d) cultures supplied with (b, e) or without (a, d) 200 µmol-N L-arginine. Abiotic control with L-arginine was also monitored (c). In a separate experimental, conversion of ^15^NO_2_^−^ to ^15^N-labeled N_2_O in the M2W compost was examined under oxic condition (f). Data are shown as means ± standard deviations (*n* = 3).

Second, to determine whether regenerated NO_2_⁻ could serve as a precursor for N_2_O under oxic conditions, M2W compost suspensions were incubated with ^15^NO_2_⁻ (Fig. 5f). Of the 1 mM ^15^NO_2_^−^ supplied, 0.016±0.003 mM and 0.698±0.065 mM were recovered as ^45^N_2_O-N and ^46^N_2_O-N, respectively. Neither sterilized controls nor incubations lacking ^15^NO_2_⁻ produced detectable labeled N_2_O. Together, these physiological experiments demonstrate that L-arginine metabolism can regenerate oxidized nitrogen, which subsequently serves as a precursor for aerobic N_2_O production independently of canonical ammonia oxidation.

### Compost metagenomes encode extensive denitrification capacity

Metagenomic analyses further demonstrated substantial genetic potential for denitrification across all compost samples (Fig. 4; Supplementary Table 6). Among genes encoding dissimilatory nitrate reductases, *narG* predominated throughout composting, with *Enteractinococcus*-affiliated clusters dominating the MI and M2W samples and *Bacilli*-affiliated clusters dominating the CC sample. Nitrite reduction potential was similarly abundant, with an *Ornithinibacillus*-affiliated *nirK* cluster representing the dominant NO-forming nitrite reductase in the MI and M2W samples.

Genes encoding nitric oxide and nitrous oxide reduction (*norB* and particularly *nosZ*) were consistently less abundant than those encoding nitrate and nitrite reduction. The M2W sample, which exhibited the highest N_2_O production under oxygen-limited conditions, showed especially low *nosZ* abundance (0.03 copies/*rpoB*) relative to *nirK* (0.32 copies/*rpoB*) and *norB* (0.004 copies/*rpoB*). Majority of *nosZ* sequences belonged to Bacilli (clade II), and qNor genes predominated over cNor genes in the UM, MI, and M2W samples. Collectively, these metagenomic data indicate that the compost microbiomes possessed extensive capacity for nitrate and nitrite reduction but comparatively limited potential for complete denitrification, particularly N_2_O reduction. These genomic profiles were consistent with the pronounced N_2_O accumulation observed during incubation.

## Discussion

Our results demonstrate that substantial N_2_O production can arise from organic nitrogen during poultry manure composting in the apparent absence of canonical nitrification. Independent evidence from inhibitor experiments, 16S rRNA gene profiling, and shotgun metagenomics consistently indicated that ammonia-oxidizing microorganisms contributed little, if at all, to nitrogen turnover in the investigated composts. Instead, our observations support an alternative conceptual model in which NO_2_^−^ and NO_3_^−^ are generated independently of ammonia oxidation and subsequently converted to N_2_O through denitrification. Although several components of this pathway remain inferential, the combined physiological, genomic, and isotope-based evidence suggests that bacterial nitric oxide synthase (bNOS)-mediated nitric oxide formation coupled with aerotolerant denitrification may represent a previously overlooked route of N_2_O production in composting systems, characterized by high temperature and organic contents.

A central question arising from our observations was the origin of NO_2_^−^ and NO_3_^−^ despite the apparent absence of canonical nitrifiers. Under conventional models, accumulation of oxidized nitrogen in manure would be interpreted as evidence of ammonia oxidation^3, 4, 5, 6^. However, neither *amoA* nor recognizable ammonia-oxidizing archaea or bacteria were detected in sequence analyses, and ATU failed to suppress N_2_O production or alter nitrogen transformations. Instead, metagenomic analyses identified abundant *bNos* genes throughout the composting process. Together with the observation that *Bacillus* isolates carrying bNOS transiently produced NO_2_⁻ and NO_3_⁻ from L-arginine, these findings suggest that nitric oxide generated from organic nitrogen may constitute an alternative source of oxidized nitrogen^11, 12, 13, 14^. Moreover, the rapid abiotic oxidation of NO by O_2_ provides a chemically plausible mechanism for the simultaneous formation of both NO_2_⁻ and NO_3_⁻^15^. Overall, this mechanism provides a coherent explanation for the accumulation of oxidized nitrogen despite the apparent absence of canonical ammonia oxidation in the compost microbiome.

Although direct *in situ* activity of bNOS remains to be demonstrated, several observations support its potential ecological relevance. Bacilli, which include many bNOS-encoding bacteria, dominated the microbial communities throughout composting, and bNOS genes affiliated with *Bacillus* spp. were consistently recovered from all metagenomes^16, 17, 18^. The accumulation of NO_2_⁻ and NO_3_⁻ during nominally anoxic incubations may seem inconsistent with the oxygen requirement of bNOS activity and abiotic NO oxidation^12, 18^. However, compost matrices are intrinsically heterogeneous and likely retain localized O_2_ as diffusion limitations generate steep O_2_ gradients despite N_2_ flushing^8, 10^. Such microoxic environments could sustain localized NO production by bNOS and subsequent oxidation, while adjacent oxygen-limited regions remain favorable for denitrification. Thus, NO_2_^−^/NO_3_^−^ formation from organic nitrogen and denitrification need not occur as spatially separated aerobic and anaerobic processes but may instead be tightly coupled across adjacent microenvironments within the same compost matrix.

Our results further identify denitrification as the primary pathway converting NO_2_^−^ and NO_3_^−^ to N_2_O. Recovery of ^15^N-labelled N_2_O following ^15^NO_2_^−^ addition demonstrated that NO_2_^−^ served as an immediate precursor of emitted N_2_O under aerobic conditions, whereas the absence of labelled N_2_O in sterilized controls excluded abiotic reduction. Continued accumulation of labelled N_2_O is consistent with differential oxygen sensitivities among sequential denitrification enzymes, with nitrous oxide reductase generally being the most oxygen-sensitive^19, 20, 21, 22, 23^. Metagenomic analyses provided a complementary explanation for the observed N_2_O accumulation. Genes encoding nitrate and nitrite reduction remained abundant throughout composting processes, whereas *nosZ* was comparatively underrepresented, particularly in the M2W samples exhibiting the highest N_2_O emissions^24, 25, 26, 27^. Phylogenetic reconstruction further indicated that many of the dominant organisms were modular denitrifiers capable of producing NO and N_2_O, but lacking the genetic capacity for N_2_O reduction. Such community organization may be particularly advantageous in thermophilic manure environments, where N_2_O reductase is known to be more temperature sensitive than upstream denitrification enzymes^28^. These observations suggest that elevated N_2_O emissions need not arise solely from transient physiological inhibition of nitrous oxide reductase but may also reflect an intrinsic imbalance in community-level denitrification capacity, whereby organisms capable of N_2_O production substantially outnumber those capable of its consumption.

Collectively, our findings support a conceptual model in which N_2_O production from organic nitrogen during poultry manure composting is governed less by canonical nitrification than by interactions between an alternative NO-generating pathway and modular denitrification within heterogeneous compost microenvironments. This framework provides a plausible explanation for the frequent observation that substantial N_2_O emissions occur in thermophilic composts despite little or no convincing evidence for active ammonia oxidation^3, 4, 5, 6, 29^. More broadly, similar conditions - high organic nitrogen availability, elevated temperature, and microbial communities dominated by facultative heterotrophs - occur in numerous engineered and natural systems, including other manure composts and potentially anaerobic digesters^30, 31, 32^. Thus, the contribution of canonical nitrification to N_2_O emissions from protein-rich thermophilic environments may warrant re-evaluation, while alternative pathways such as the one proposed here deserve greater attention in future studies aimed at understanding and mitigating biological N_2_O production.

## Acknowledgements

This work was financially supported by the National Research Foundation of Korea (NRF) grant funded by the Korean government (RS-2026-25525273). MS was also supported by an NRF grant funded by the Korean government (MSIT) (RS-2024-00337570).

## Author information

### Contributions

SY conceived and supervised the study. JYL and MJS designed experiments. JYL performed experiments and analyzed experimental and sequencing data. MJS and ML supported data analysis. JYL and SY wrote manuscript with contributions from MJS and ML. All authors critically reviewed the manuscript.

## Supporting Information

Supplemental information text (PDF)

Supplementary Tables (XLSX)

## Supporting Information

### Supplementary figures

**Supplementary Fig. 1.**
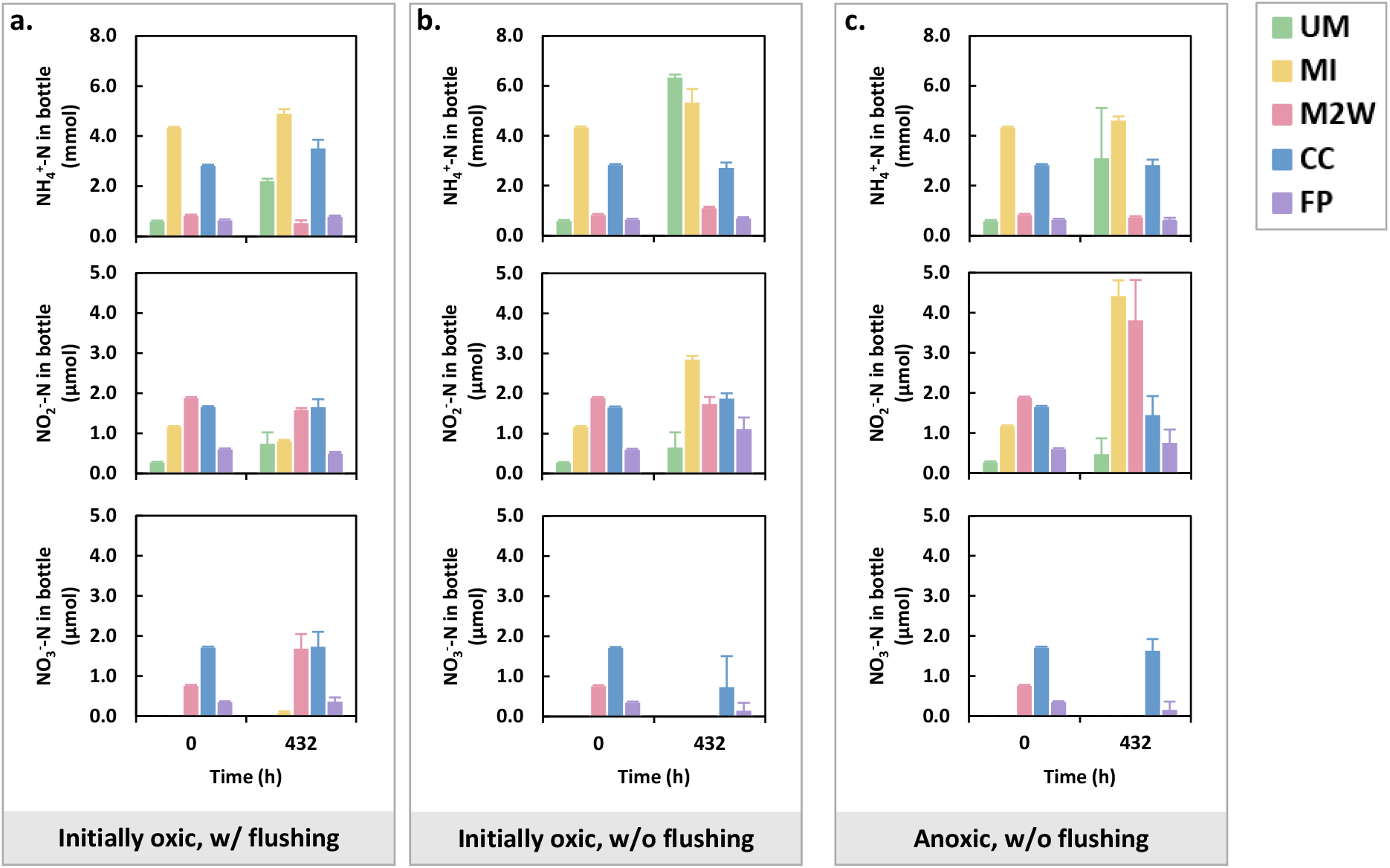
Additional data supporting Figure 1. Two-point measurements of inorganic nitrogen (NH_4_^+^-N, NO_2_ ^−^-N, and NO_3_^−^-N) concentrations in poultry manure and compost samples at the beginning and end of incubations. Values represent the means of triplicate incubations, and the error bars indicate standard deviations.

## Reference

1. Wuebbles, D. J. Nitrous Oxide: No laughing matter. Science 326, 56–57 (2009).

2. Prosser, J. I., Hink, L., Gubry-Rangin, C. & Nicol, G. W. Nitrous oxide production by ammonia oxidizers: Physiological diversity, niche differentiation and potential mitigation strategies. Global Change Biology 26, 103–118 (2020).

3. Xu, Z., et al. Microbial sources and sinks of nitrous oxide during organic waste composting. Environmental Science & Technology 58, 7367–7379 (2024).

4. Fukumoto, Y., et al. Reduction of nitrous oxide emission from pig manure composting by addition of nitrite-oxidizing bacteria. Environmental Science & Technology 40, 6787–6791 (2006).

5. He, Y., et al. Nitrous oxide emissions from aerated composting of organic waste. Environmental Science & Technology 35, 2347–2351 (2001).

6. Maeda, K., et al. Microbiology of nitrogen cycle in animal manure compost. Microbial Biotechnology 4, 700–709 (2011).

7. Hou, Y., Velthof, G. L., Lesschen, J. P., Staritsky, I. G. & Oenema, O. Nutrient recovery and emissions of ammonia, nitrous oxide, and methane from animal manure in Europe: Effects of manure treatment technologies. Environmental Science & Technology 51, 375–383 (2017).

8. Nordahl, S. L., Preble, C. V., Kirchstetter, T. W. & Scown, C. D. Greenhouse gas and air pollutant emissions from composting. Environmental Science & Technology 57, 2235–2247 (2023).

9. Zumft, W. G. Cell biology and molecular basis of denitrification. Microbiology and Molecular Biology Reviews 61, 533–616 (1997).

10. Szanto, G. L., Hamelers, H. V. M., Rulkens, W. H. & Veeken, A. H. M. NH_3_, N_2_O and CH_4_ emissions during passively aerated composting of straw-rich pig manure. Bioresource Technology 98, 2659–2670 (2007).

11. Adak, S., Aulak, K. S. & Stuehr, D. J. Direct evidence for nitric oxide production by a nitric-oxide synthase-like protein from *Bacillus subtilis*. Journal of Biological Chemistry 277, 16167–16171 (2002).

12. Gusarov, I., et al. Bacterial nitric-oxide synthases operate without a dedicated redox partner. Journal of Biological Chemistry 283, 13140–13147 (2008).

13. van Sorge, N. M., et al. Methicillin-resistant *Staphylococcus aureus* bacterial nitric-oxide synthase affects antibiotic sensitivity and skin abscess development. Journal of Biological Chemistry 288, 6417–6426 (2013).

14. Wang, Z.-Q., et al. Bacterial flavodoxins support nitric oxide production by *Bacillus subtilis* nitric-oxide synthase. Journal of Biological Chemistry 282, 2196–2202 (2007).

15. Ignarro, L. J., Fukuto, J. M., Griscavage, J. M., Rogers, N. E. & Byrns, R. E. Oxidation of nitric oxide in aqueous solution to nitrite but not nitrate: comparison with enzymatically formed nitric oxide from L-arginine. Proceedings of the National Academy of Sciences 90, 8103–8107 (1993).

16. Arora, D. P., Hossain, S., Xu, Y. & Boon, E. M. Nitric oxide regulation of bacterial biofilms. Biochemistry 54, 3717–3728 (2015).

17. Gusarov, I., Shatalin, K., Starodubtseva, M. & Nudler, E. Endogenous nitric oxide protects bacteria against a wide spectrum of antibiotics. Science 325, 1380–1384 (2009).

18. Sudhamsu, J. & Crane, B. R. Bacterial nitric oxide synthases: what are they good for? Trends in Microbiology 17, 212–218 (2009).

19. Betlach, M. R. & Tiedje, J. M. Kinetic explanation for accumulation of nitrite, nitric oxide, and nitrous oxide during bacterial denitrification. Applied and Environmental Microbiology 42, 1074–1084 (1981).

20. Körner, H. & Zumft, W. G. Expression of denitrification enzymes in response to the dissolved oxygen level and respiratory substrate in continuous culture of *Pseudomonas stutzeri*. Applied and Environmental Microbiology 55, 1670–1676 (1989).

21. Morley, N., Baggs, E. M., Dörsch, P. & Bakken, L. Production of NO, N_2_O and N_2_ by extracted soil bacteria, regulation by NO_2_^−^ and O_2_ concentrations. FEMS Microbiology Ecology 65, 102–112 (2008).

22. Roothans, N., et al. Aerobic denitrification as an N_2_O source from microbial communities. The ISME Journal 18, 1 (2024).

23. Tang, W., et al. Similar oxygen sensitivities of different steps of denitrification in Estuarine waters. Environmental Science & Technology 59, 7165–7175 (2025).

24. Castellano-Hinojosa, A., González-López, J. & Bedmar, E. J. Distinct effect of nitrogen fertilisation and soil depth on nitrous oxide emissions and nitrifiers and denitrifiers abundance. Biology and Fertility of Soils 54, 829–840 (2018).

25. Henry, S., Bru, D., Stres, B., Hallet, S. & Philippot, L. Quantitative detection of the *nosZ* gene, encoding nitrous oxide reductase, and comparison of the abundances of 16S rRNA, *narG*, *nirK*, and *nosZ* genes in soils. Applied and Environmental Microbiology 72, 5181–5189 (2006).

26. Pessi, I. S., et al. In-depth characterization of denitrifier communities across different soil ecosystems in the tundra. Environmental Microbiome 17, 30 (2022).

27. Roothans, N., et al. Long-term multi-meta-omics resolves the ecophysiological controls of seasonal N_2_O emissions during wastewater treatment. Nature Water 3, 590–604 (2025).

28. Ma, L., et al. Temperature-dependent regulation of denitrification intermediates in high-temperature ecosystems. Environmental Science & Technology 59, 12630–12641 (2025).

29. Cáceres, R., Malińska, K. & Marfà, O. Nitrification within composting: A review. Waste Management 72, 119–137 (2018).

30. Beffa, T., et al. Taxonomic and metabolic microbial diversity during composting. In: The Science of Composting (eds de Bertoldi M., Sequi P., Lemmes B., Papi T.). Springer Netherlands (1996).

31. Finore, I., et al. Thermophilic bacteria and their thermozymes in composting processes: a review. Chemical and Biological Technologies in Agriculture 10, 7 (2023).

32. Westerholm, M., et al. Microbial community dynamics linked to enhanced substrate availability and biogas production of electrokinetically pre-treated waste activated sludge. Bioresource Technology 218, 761–770 (2016).

33. Ginestet, P., Audic, J.-M., Urbain, V. & Block, J.-C. Estimation of nitrifying bacterial activities by measuring oxygen uptake in the presence of the metabolic inhibitors allylthiourea and azide. Applied and Environmental Microbiology 64, 2266–2268 (1998).

34. Yoshinari, T., Hynes, R. & Knowles, R. Acetylene inhibition of nitrous oxide reduction and measurement of denitrification and nitrogen fixation in soil. Soil Biology and Biochemistry 9, 177–183 (1977).

35. Bolyen, E., et al. Reproducible, interactive, scalable and extensible microbiome data science using QIIME 2. Nature Biotechnology 37, 852–857 (2019).

36. Bolger, A. M., Lohse, M. & Usadel, B. Trimmomatic: a flexible trimmer for Illumina sequence data. Bioinformatics 30, 2114–2120 (2014).

37. Callahan, B. J., et al. DADA2: High-resolution sample inference from Illumina amplicon data. Nature Methods 13, 581–583 (2016).

38. Robeson, M. S., II, et al. RESCRIPt: Reproducible sequence taxonomy reference database management. PLOS Computational Biology 17, e1009581 (2021).

39. Lee, J. Y., et al. Selective enrichment of methylococcaceae versus methylocystaceae methanotrophs via xontrol methane feeding schemes. Environmental Science & Technology 58, 14237–14248 (2024).

40. Chen, S., Zhou, Y., Chen, Y. & Gu, J. fastp: an ultra-fast all-in-one FASTQ preprocessor. Bioinformatics 34, i884–i890 (2018).

41. Nurk, S., Meleshko, D., Korobeynikov, A. & Pevzner, P. A. metaSPAdes: a new versatile metagenomic assembler. Genome Res 27, 824–834 (2017).

42. Hyatt, D., et al. Prodigal: prokaryotic gene recognition and translation initiation site identification. BMC Bioinformatics 11, 119 (2010).

43. Aramaki, T., et al. KofamKOALA: KEGG Ortholog assignment based on profile HMM and adaptive score threshold. Bioinformatics 36, 2251–2252 (2019).

44. Buchfink, B., Xie, C. & Huson, D. H. Fast and sensitive protein alignment using DIAMOND. Nature Methods 12, 59–60 (2015).

45. Murali, R., et al. Diversity and evolution of nitric oxide reduction in bacteria and archaea. Proceedings of the National Academy of Sciences 121, e2316422121 (2024).

46. Jones, P., et al. InterProScan 5: genome-scale protein function classification. Bioinformatics 30, 1236–1240 (2014).

47. Li, H. Aligning sequence reads, clone sequences and assembly contigs with BWA-MEM. arXiv: Genomics (2013).

48. Li, H., et al. The Sequence alignment/map format and SAMtools. Bioinformatics 25, 2078–2079 (2009).

49. Quinlan, A. R. & Hall, I. M. BEDTools: a flexible suite of utilities for comparing genomic features. Bioinformatics 26, 841–842 (2010).

50. Kjeldahl, J. Neue methode zur bestimmung des stickstoffs in organischen Körpern. Zeitschrift für analytische Chemie 22, 366–382 (1883).

51. Walkley, A. A Critical examination of a rapid method for determining organic carbon in soil—Effect of variations in digestion conditions and of inorganic soil constituents. Soil Science 63, (1947).

52. Chang, J., et al. Enhancement of nitrous oxide emissions in soil microbial consortia via copper competition between proteobacterial methanotrophs and denitrifiers. Applied and Environmental Microbiology 87, e02301–02320 (2021).

53. Miranda, K. M., Espey, M. G. & Wink, D. A. A rapid, simple spectrophotometric method for simultaneous detection of nitrate and nitrite. Nitric Oxide 5, 62–71 (2001).

54. Baethgen, W. E. & Alley, M. M. A manual colorimetric procedure for measuring ammonium nitrogen in soil and plant Kjeldahl digests. Communications in Soil Science and Plant Analysis 20, (1989).

55. Wentworth, W. E. & Freeman, R. R. Measurement of atmospheric nitrous oxide using an electron capture detector in conjunction with gas chromatography. Journal of Chromatography A 79, 322–324 (1973).

56. Yoon, S., Nissen, S., Park, D., Sanford, R. A. & Löffler, F. E. Nitrous oxide reduction kinetics distinguish bacteria harboring clade I NosZ from those harboring clade II NosZ. Applied and Environmental Microbiology 82, 3793–3800 (2016).

57. Kuang, C., et al. Accurate quantification of ^45^N_2_O with a membrane inlet mass spectrometer by optimizing the temperature of a cold trap. JACS Au 5, 5821–5827 (2025).

58. Sander, R. Compilation of Henry’s law constants (version 4.0) for water as solvent. Atmospheric Chemistry and Physics 15, 4399–4981 (2015).

59. Ridwan, R., Abdul Razak, H. R., Adenan, M. I. & Md Saad, W. M. Development of isocratic RP-HPLC method for separation and quantification of L-citrulline and L-arginine in watermelons. International Journal of Analytical Chemistry 2018, 4798530 (2018).

